# Pupil Dynamics reflect Compensatory Control during Continuous Motor-Control following Cognitive Demand

**DOI:** 10.64898/2026.07.30.741925

**Authors:** Esther A. Semmelhack, Anne Schacht

## Abstract

Pupillometry is commonly used to assess workload and fatigue, but its interpretation in applied, continuous-control tasks remains unclear because pupil responses are strongly shaped by task context and often lack a theory-guided interpretive basis. The Compensatory Control Model and Locus Coeruleus-Norepinephrine system offer complementary frameworks for interpreting pupil dynamics in relation to regulatory effort and engagement under sustained cognitive demand. We aimed to examine how tonic and phasic pupil dynamics reflect compensatory regulation following sustained cognitive demand, and how these dynamics relate to subsequent performance during continuous motor control. Participants completed two sessions consisting of a low- versus high-demand working-memory task followed by a joystick-based continuous motor-control task. Linear mixed-effects models tested demand effects on pupil dynamics and behavior and evaluated trial-level pupil–performance associations. Prior high demand produced small and specific changes in motor-control performance. Task-evoked pupil response showed effects of engagement dependent on prior demand and small associations with movement timing and accuracy. Our results show that pupil dynamics sensitively reflect sustained cognitive demand, but their trial-level predictive value for subsequent continuous motor performance is limited.

## Introduction

In safety-critical settings, such as vehicle control or human-robot-collaboration (HRC), changes in operator internal state fluctuation can affect performance and thereby compromise safety (1–4). Accordingly, driver state monitoring systems are used to track indicators of reduced alertness and drowsiness, often based on vision-derived eye measures (5–7). In HRC, safety approaches are commonly framed around physical safeguards, whereas evidence for continuous psychophysiological state monitoring in HRC remains limited (4,8–11). Current user monitoring in HRI focuses largely on coarse performance metrics like response time (RT) and accuracy (10). Despite their relevance for collaborative actions such as handovers, fine-grained motor-control measures, such as movement precision, remain underrepresented in HRC research (10,11). Moreover, physiological indicators have been explored for performance estimation in HRI, but existing approaches often lack a clear theoretical account of which operator states are indexed and how systems should respond to them (10,12,13). Misinterpreting physiological signals may lead to inappropriate system responses in safety-critical interaction contexts (3,13,14). Taken together, these findings motivate the need for physiological indicators that can complement performance-based monitoring by capturing fluctuations in operator state during continuous interaction, as well as for a clear conceptual framework to interpret such signals appropriately in safety-critical HRC contexts.

Pupil diameter and pupil dynamics are promising candidates for operator-state monitoring, as they can be measured remotely via infrared cameras and have been linked to arousal, effort, and engagement-related states (12,15–17). In controlled settings, pupil diameter and other physiological indicators are often sensitive to workload manipulations and show modest associations with performance, but these relationships can vary substantially across individuals and task contexts, underscoring the need for validation before applied monitoring (18). Psychometric evidence further indicates that physiological workload measures are often only weakly related to one another and may reflect distinct underlying processes rather than a single unitary “workload” signal (19). Evidence for these relationships has been established primarily in controlled laboratory paradigms with short- duration tasks and discrete responses, which yield reliable effects but raise questions about their transferability to continuous, HRC-relevant control contexts (10,20). Applied HRC research has begun to incorporate pupillometry in collaborative scenarios, for example, to examine workload or learning-related processes, but this literature is often based on limited samples or descriptive analyses, constraining theory-driven interpretation (11,21). Recent work has further explored pupil measures for estimating or predicting task performance, alongside modeling approaches that address uncertainty in human-state inference (12,13). However, pupil dynamics are highly context dependent, varying with task difficulty, emotion, motivation, attentional fluctuations, and mind wandering, indicating that similar pupil patterns may correspond to different underlying states (22–25). For adaptive systems, this interpretive ambiguity is consequential because similar physiological patterns may reflect different underlying states or resource demands, requiring careful, theory-guided interpretation to avoid mis-targeted or inefficient system responses (13,19). Together, these considerations motivate theory-guided studies of pupil dynamics that balance experimental control with ecologically relevant interaction demands, and that support interpretable and context-appropriate use of physiological indicators in HRC.

To interpret pupil dynamics in a meaningful way, we draw on two complementary frameworks: the Compensatory Control Model (CCM; Hockey, 1997, 2011) explains how people maintain performance through effort regulation, and the locus coeruleus-norepinephrine (LC-NE) system specifies how these regulatory processes are expressed physiologically in tonic and phasic pupil activity. According to the CCM people can maintain performance under sustained or increasing cognitive demand by regulating effort and adjusting their control strategies: Performance can remain relatively stable under elevated demand even as internal strain increases, because additional effort is recruited to compensate for rising costs. However, compensatory regulation is limited, and prolonged demand ultimately leads to strategic shifts or performance deterioration (26,27). Hence, pupil dynamics may provide complementary information about fluctuations in effort and engagement during continuous perceptual–motor control. Accordingly, the present study conceptualizes operator state as a continuous and dynamically fluctuating process of compensatory regulation, reflected in ongoing adjustments of behavior and pupil dynamics. Specifically, we focus on compensatory control following sustained cognitive demand, on the assumption that regulatory states induced by prolonged effort can persist across task boundaries and shape subsequent performance during continuous control.

Models of the locus coeruleus–norepinephrine (LC–NE) system provide a physiological framework for interpreting pupil dynamics in relation to arousal and task engagement (16,28–30). Non-luminance-mediated pupil changes are linked to moment-to-moment LC–NE activity: tonic LC– NE activity reflects global arousal and motivational state and is associated with baseline pupil diameter, whereas phasic LC–NE bursts support selective attention and task engagement and are associated with task-evoked dilations. Consistent with this framework, task-evoked (phasic) pupil responses consistently decrease with disengagement, attentional lapses, or prolonged task engagement (31–33). Importantly, reduced task-evoked pupil responses can be restored by increasing reward, indicating sensitivity to motivational control rather than irreversible resource depletion (31). In contrast, baseline pupil diameter shows inconsistent patterns across studies. Decreases in baseline pupil diameter have been reported under prolonged effort, fatigue, or sleepiness (34), whereas increases have been associated with mind-wandering, exploratory states, or disengagement in reading and vigilance tasks (24,25). Other studies report mixed or state-dependent tonic effects: baseline pupil diameter did not consistently track performance decline over time (33), and different types of attentional lapses were associated with opposite baseline patterns despite similarly reduced phasic responses (32). Together, these findings indicate that tonic pupil diameter reflects broad arousal and motivational context rather than a specific form of cognitive demand or fatigue, and that pupil–state relationships are strongly shaped by task structure and context. Accordingly, pupil dynamics are best interpreted as reflecting continuous, context-dependent regulation of cognitive control rather than discrete or categorical internal states. Together, CCM and LC–NE offer complementary accounts of compensatory effort and its physiological expression, providing a basis for interpreting tonic and phasic pupil dynamics in sustained-demand tasks.

## Objective

Building on the need for more context-sensitive indicators of operator state, the present study examines how tonic and phasic pupil dynamics relate to continuous hand–eye motor-control after sustained cognitive demand in a joystick-based task inspired by HRC. More specifically, we aim to characterize distinct components of the pupil response and its covariation with behavioral adjustments during continuous control. By integrating the Compensatory Control Model and the LC– NE adaptive gain framework, we interpret tonic and phasic pupil measures in relation to compensatory regulation. Rather than manipulating cognitive demand during the motor-control task itself, the study focuses on how sustained prior demand influences subsequent behavioral regulation and pupil dynamics, thereby dissociating regulatory state from momentary task load.

Participants completed two working-memory tasks that differed in cognitive demand and duration before performing the joystick-based motor-control task. Comparisons between low- and high-demand tasks are investigated only for validation of cognitive demand. For the subquent motor task, this study tests three hypotheses:

### Hypothesis 1

Motor performance during continuous control will show selective differences as a function of prior cognitive demand, indicating carry-over effects of compensatory regulation.

### Hypothesis 2

Prior cognitive demand will be reflected in pupil dynamics during the motor-control task, with phasic, task-evoked pupil responses showing greater sensitivity to demand-related differences than tonic baseline pupil diameter

### Hypothesis 3

Pupil responses will covary with moment-to-moment fluctuations in motor performance.

## Method

### Transparency and Openness

The study was pre-registered prior to data collection (https://osf.io/udmv9/). We report sample-size determination, all data exclusions, manipulations, and measures. All preprocessed data are openly available on the Open Science Framework (https://osf.io/6ypuf/files/osfstorage). Full methods details are provided in the Supplementary Information.

### Participants

For primary sample size estimation, we used GPower for a within-subject low vs. high demand contrast, basing the effect size on a pupillometry effect from Unsworth et al. (2018), chosen as a conservative proxy because pupil effects are typically smaller than behavioral endpoints in our paradigm, rather than generic ‘small/medium/large’ conventions (Correll et al., 2020). GPower inputs were: t-test, Means: difference between dependent samples, two-tailed, dz=0.428, power = 0.80, α=.05. The resulting target was N=45 (noncentrality = 2.87, critical t = 2.02, df = 44, actual power = 0.801). GPower was used only to size the primary within-subject contrast and is not applicable to interaction terms or random-slope structures in our mixed models. Throughout the manuscript, we report semi-partial R² for fixed effects because it provides a term-level, scale-free index appropriate for linear mixed models (i.e., it reflects the unique variance attributable to each predictor and is comparable across outcomes).

Forty-six participants were recruited (24 male, 23 female); of these, 40 (18 male, 22 female) completed both sessions. Data from all available sessions were used. Age ranged between 19 and 35 years (M=22.95, SD=3.26). All were either native German speakers or had proficient German or English, were students at the Georg-August-University of Göttingen, had normal or corrected-to- normal vision (+/–1 diopter), no neurological or psychiatric history, and no joystick experience. Participants received either course credits or 10 Euros per hour as compensation. Recruitment started 07.01.2024 and ended 23.07.2024, data collection started 22.01.2024 and was completed 30.07.204.

This research complied with the American Psychological Association Code of Ethics and was approved by the Ethics Committee of the Institute of Psychology at the University of Göttingen. Informed, written consent was obtained from each participant.

### Design and Procedure

All participants were tested individually in a sound-attenuated EEG cabin. They were seated on an adjustable chair and used a chinrest at a fixed viewing distance of 60 cm from a 24-inch monitor. Participants were instructed to maintain central fixation and minimize blinking throughout all task blocks to ensure data quality, particularly for pupil recordings. The experiment consisted of two sessions of approximately 2.5 hours each, spaced exactly one week apart. Session order was counterbalanced such that half of the participants began with the low-demand task and the other half with the high-demand task. The one-week interval was chosen to maximize comparability in baseline mental state due to similar weekly schedules.

Each session began with either the low- or high-demand task (order counterbalanced), followed by the motor-control task. At the beginning of the first session, participants completed a brief demographic questionnaire, and filled out the Stanford Sleepiness Scale (35), the PANAS (36), and a 10-point visual fatigue scale (37). Practice trials were completed before each task to ensure understanding. Eye tracking and EEG were recorded throughout the sessions. EEG data, mood and sleepiness scores were not central for the current research aims and are not reported here.

In the low-demand condition, participants completed a short two-back task (38). Eight different uppercase letters were presented in random order at the center of the screen. Participants were instructed to press the space bar whenever the current letter matched the one presented two trials earlier. Practice trials were completed first to ensure task understanding. Each trial began with a 500 ms fixation cross, followed by a 1200 ms stimulus presentation. Thirty percent of trials were two-back targets, and no more than two occurred consecutively. The same letter was never repeated more than twice in a row. The task was divided into four blocks, each consisting of 32 trials (eight letters presented four times). Participants were required to take a 30-second break between blocks and resume as soon as the time expired.

In the high-demand condition, participants completed the time-load dual-back (TLDB) task (39). Stimuli alternated between uppercase letters and single-digit numbers. Participants used both hands: one for responding to two-back letter matches, and the other to categorize numbers as odd or even. Hand assignments were counterbalanced. As in the low-demand task, practice blocks preceded the main task. Each trial began with a 500 ms fixation cross, followed by a 1200 ms stimulus. Thirty percent of trials were targets (two-back or odd/even), with no more than two consecutive targets. No stimulus appeared more than twice in a row, and odd/even frequencies were balanced. The TLDB task consisted of 10 blocks of 64 stimuli (16 unique items repeated four times). If accuracy in a block exceeded 85%, stimulus duration in the next block was shortened by 150 ms; otherwise, it was increased by 150 ms. Each block was followed by a mandatory 30-second break.

After each demand task, participants again rated their mental fatigue and mood. This was followed by the motor-control task, a joystick-controlled drag-and-drop task. Participants used a Thrustmaster TCA Sidestick Airbus Edition joystick, which was individually positioned for comfort and suitable for both left- and right-handed participants. Again, practice trials were conducted. Each trial began with a 500 ms blank screen, followed by a 4500 ms fixation cross to allow pupil size to return to baseline. During this time, participants were instructed to release pressure on the joystick to allow it to recenter, and to maintain central fixation. The task display then showed three elements simultaneously: (1) a red object, (2) a robot hand in one of eight fixed locations at the screen periphery, and (3) a cursor at the screen center. The red object appeared at a fixed distance from the robot hand; the cursor always began in the center, equidistant from the object. Participants were instructed to grab the object by pressing and holding a joystick button, drag it to the robot hand, and release it accurately. Trial duration varied based on movement speed. The task included five blocks of 32 trials each – with all 8 robot and corresponding object positions being presented 20 times, for a total of 160 trials.

After the motor-control task, participants completed the Engagement Questionnaire (40), and rated their mental fatigue and mood. Participants were then debriefed. Figure 1 shows the full tsk and trial structure for both experimental sessions.

**Fig 1.**
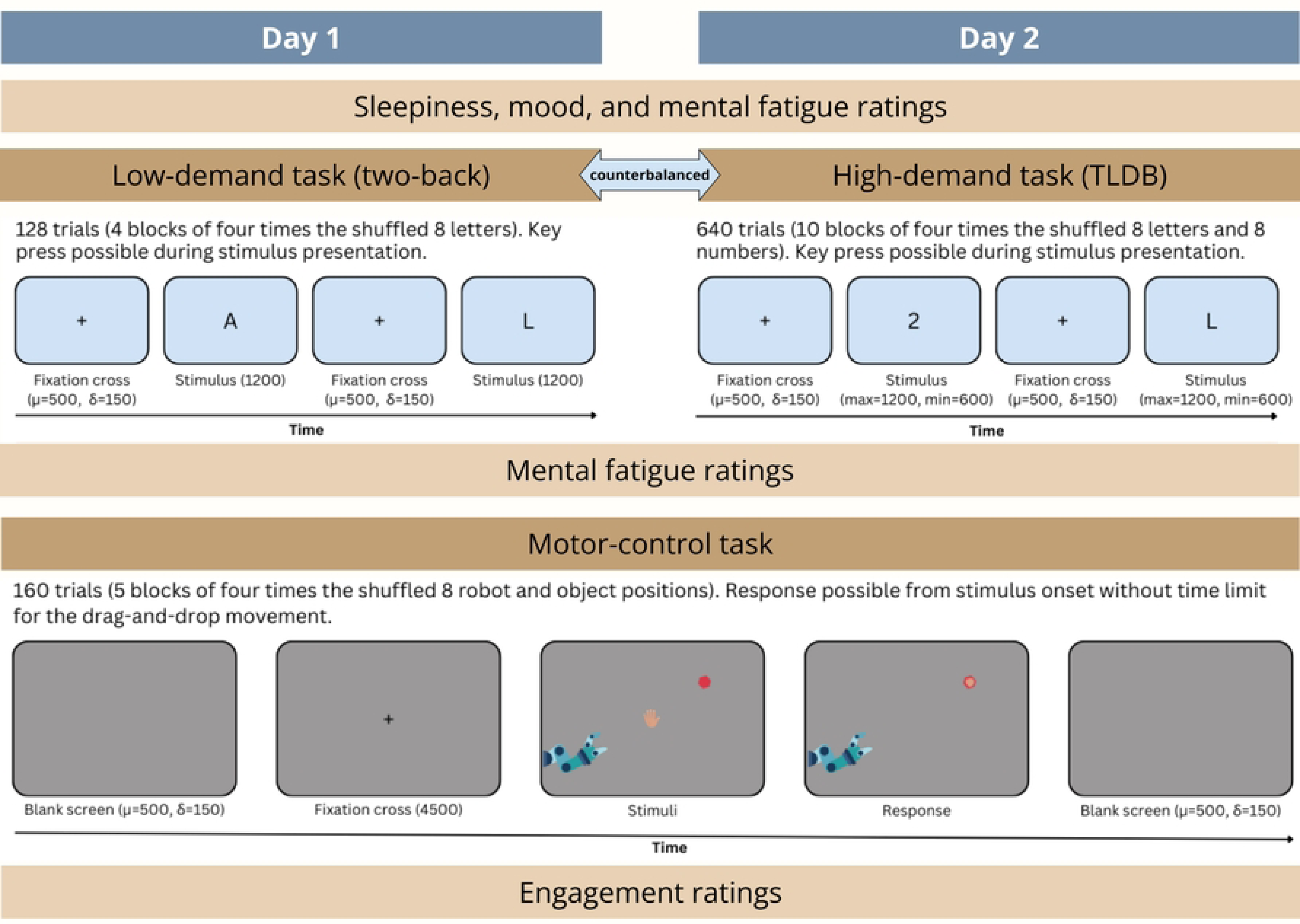
Schematic overview of the experimental procedure and trial structure.

### Apparatus and Stimuli

All tasks were presented using PsychoPy (v2023.2.3., Peirce et al., 2019) and Pygame (v2.5.2., Shinners, 2011) with joystick input handled via PyLink (v2.1.762.0). Stimuli were presented on a 24- inch LCD monitor (1920 × 1080 pixels, 60 Hz refresh rate). The motor-control task used a Thrustmaster TCA Sidestick Airbus Edition joystick. Eye tracking was recorded with a desktop-mounted EyeLink 1000 Plus system (SR Research).

Visual stimuli for the two-back and time-load dual-back (TLDB) tasks were centrally presented in black Open Sans font on a light gray background. The stimuli consisted of eight uppercase letters from the German alphabet (A, C, D, H, L, N, T, U). Letters were selected based on high visual discriminability and comparable frequency of occurrence in both German and English, using data from the *Zeichen- und Buchstabenhäufigkeitsliste* of the Institut für Deutsche Sprache in Mannheim (43). Letters visually similar to digits (e.g., I) were excluded to avoid confusion. In the two-back task, only these letters were presented in randomized order to familiarize participants with the experimental procedure while avoiding under-arousal. In the TLDB task, the same letter set was used alongside eight single-digit numbers (two to nine), balanced for even and odd frequency. Digits visually confusable with letters (e.g., 1) were excluded. Letter and number stimuli were presented in alternating sequence.

The motor-control task was used to assess behavioral and physiological effects of the preceding cognitive demand condition (low or high). Visual stimuli in the motor control task consisted of a robotic arm appearing in one of eight different positions along the screen edges, a red hexagonal object, and cartoon-style hand serving as the cursor (see Figure 1). Stimuli were designed using Procreate for IPad (version 5.3.4) and converted to scalable vector graphics using GIMP (GNU Image Manipulation Program, version 2.10.34). All stimuli were custom-made for this experiment and have not been used in previous studies.

### Pupil recording and pre-processing

Pupil size was recorded binocularly in arbitrary units (AUs) at 500 Hz, following a nine-point calibration and validation procedure. A fake-pupil recording was used to transform pupil size data from arbitrary units to millimeter.

Preprocessing of pupil data was performed in MATLAB (R2018a) following Kret and Sjak-Shie (44). Pupil time series were time-locked to the onset of the target stimuli in both the demand task (letters and numbers) and the motor-control task (robot arm, object, and cursor). Data samples were classified as blinks or invalid gaze when data from both eyes were lost. Artifacts were removed using a multi-step procedure: Speed outlier rejection was performed by flagging samples exceeding 16 times the median absolute deviation (MAD) of pupil dilation speed as outliers. To avoid interpolation artifacts, 50 ms (25 samples) were trimmed from both sides of each data gap. Linear interpolation was applied to fill remaining gaps. A trend line was computed, and clusters of samples showing strong deviation from the trend were rejected. And finally, isolated samples surrounded by missing data were discarded. The interpolated signal was divided into contiguous segments. A second-order, 4 Hz low-pass Butterworth filter was applied separately to each segment to smooth the data while avoiding edge artifacts in short segments. This piecewise filtering approach ensured that segments with insufficient length (i.e., too few samples) were not erroneously filtered. Baseline correction was performed by subtracting the mean pupil value during the -200 ms to 0 ms interval per subject and session relative to stimulus onset from each trial’s signal.

### Statistical Analysis

All analyses were conducted in R (v4.3.2). Linear mixed-effects models (LMMs) and generalized linear mixed-effects models (GLMMs) were used to estimate effects of cognitive demand condition, time, and their interaction on behavioral and pupil measures, while accounting for random effects of subject and stimulus.

To control the Type I error rate, all identifiable random slopes were included (45,46) for the grouping factors subject ID and stimulus ID. For linear models fitted via the function lmer from the package *lme4* (47), model assumptions were normally distributed and homogenous residuals and were verified through visual inspection of the QQ plot of residuals (48) and residuals plotted against fitted values (49). For non-linear models fitted via glmmTMB, model assumptions were no over- or under-dispersion and were verified via the function overdisp.test. To assess collinearity, we re-estimated each model without the interaction terms and computed the Variance Inflation Factors using the function vif from the car package (50). Confidence intervals for model estimates were obtained via parametric bootstrapping, using the function boot.lmer of the package *lme4* (N = 100 bootstraps) or the function boot.glmmtmb from the package *boot.glmmtmb* (n = 100 bootstraps).

Behavioral performance during the cognitive-demand tasks was analyzed to validate the demand manipulation using a GLMM with beta-family error distribution for accuracy and a LMM for reaction times (RT). Both models included demand condition (low vs. high), percentage of the session (continuous) and initial fatigue score (continuous) as fixed effects, along with their two-way interactions. The overall effect of demand condition, as well as its interaction with percentage of the session and initial fatigue score was evaluated using a full-null model comparison (51). The null model excluded main effects of demand condition, percentage and their interaction but was otherwise identical in structure. Statistical significance of each level of the factor and their interaction with percentage of the session and initial fatigue was tested by means of the Satterthwaite approximation (52), via the function lmer from the package lmerTest (53). All models were fitted with low demand as the reference level of the factor demand condition.

Behavioral data from the motor-control task were analyzed using four response variables: reaction time (RT), movement precision, movement duration, and hit precision. RT (in ms) was defined as the interval between stimulus onset and the object grabbing. Movement precision (in %) was computed as the average frame-wise deviation from the ideal straight-line trajectory toward the target. Movement duration (in ms) referred to the time between grabbing and releasing the object. Hit precision (in %) was defined as the percentage of spatial overlap between the object and the target area at the time of release.

To normalize distributions, movement precisions, and movement durations were log-transformed. Outliers exceeding ±2 standard deviations (SD) from each participant’s mean were excluded (54). RTs shorter than the physiological limit of 400ms and longer than 2000 ms, trials taking longer than 10000 ms, and movement durations smaller than 400 ms and longer than 5000 ms were considered implausible and removed. Trials with zero deviation in the movement precision measures – typically due to failed object-grabs –were also excluded. For hit precision, trials with zero overlap with the target area at the moment of the release were removed and outliers beyond 2 SD were removed.

GLMMs were constructed for each behavioral metric, following the same model structure and fitting procedures described for the demand task. Fixed effects included demand condition (low vs. high), trial number (continuous), and task engagement (continuous), plus their two-way interactions. Random slopes were modeled for subject ID, and stimulus ID. Significance testing and model diagnostics (full-null comparisons, residual inspection, VIFs, bootstrapped confidence intervals) were conducted as described above. For hit precision, statistical significance was tested via the function glmmTMB using a beta-family error distribution.

Pupil data were analyzed separately for the demand and the motor-control task and separately for three pupil time-windows, including the baseline pupil diameter, early pupil constriction diameter, and late pupil re-dilation diameter defined a priori based on grand-average pupil waveforms for each task (see Figure 2). For the demand task, we analyzed baseline pupil diameter (500 ms pre-stimulus), early pupil constriction (0–600 ms post-stimulus), and late pupil re- dilation (600–1200 ms post-stimulus). For the motor-control task we analyzed baseline pupil diameter 500 ms pre-stimulus), early pupil constriction (200–1200 ms), and late pupil re-dilation (1200–2200 ms). Time windows were time-locked to the onset of stimulus.

**Fig 2.**
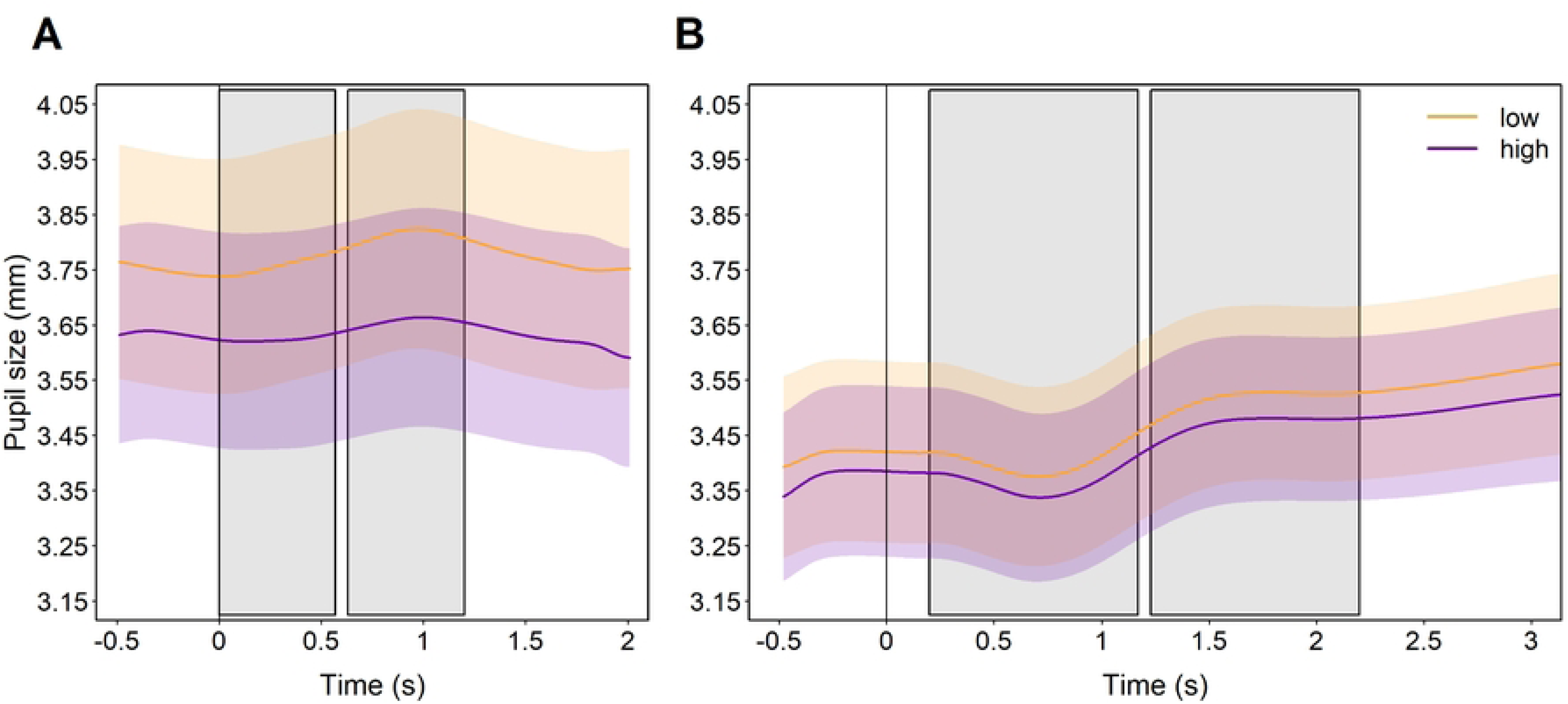
Grand-average pupil waveforms and analysis time windows. Mean pupil size for each demand condition and the corresponding analysis windows for early constriction and late re**-**dilation in (A) the cognitive demand tasks and (B) the motor-control task. Shaded areas indicate 95% confidence bands.

Prior to analysis, we applied consistent outlier exclusion criteria across both tasks and all time windows. Trial -to-trial changes in pupil size exceeding 2 SDs from each participants mean were removed. Additionally, observations outside the 2.5th and 97.5th percentiles of the overall distribution were excluded to enhance model robustness and minimize data loss.

For the cognitive demand task, fixed effects included demand condition, percentage of the session, and initial fatigue. For the motor-control task, fixed effects included demand condition, trial number, and task engagement, plus their two-way interactions. Random slopes were modeled for subject ID, and stimulus ID, except for baseline models where the stimulus was always the fixation cross, so no random slope was necessary. Significance testing and model diagnostics (full-null comparisons, residual inspection, VIFs, bootstrapped confidence intervals) were conducted as described above.

For the motor-control task, we additionally tested the predictive effect of each pupil parameter separately for each performance parameter. Fixed effects included pupil parameter (baseline, constriction, or re-dilation) in interaction with trials. Significance testing and model diagnostics were conducted as described above.

## Results

All models met assumptions of normally distributed and homoscedastic residuals and no overdispersion and revealed no collinearity issues.

### Cognitive demand task

#### Effects of demand on performance

As a manipulation check, we examined accuracy and response time (RT) in the 2-back (low demand) and TLDB (high demand) tasks.

Accuracy showed a significant interaction between demand condition and session progress: in the high-demand TLDB, accuracy declined over time, whereas in the low-demand 2-back, accuracy slightly improved *(β* = -0.408, p < .001) (Figure 3A). Accuracy also varied as a function of initial fatigue, decreasing with higher initial fatigue in the low-demand task but increasing slightly in the high-demand task (*β* = 0.304, p = .028) (Figure 3B). For RT, the interaction between demand condition and percentage of the session was significant (*β* = -0.082, p = .014, semi-partial R² = .02): RTs decreased across time in the high-demand condition, while they increased in the low-demand task (see Fig 3C).

**Fig 3.**
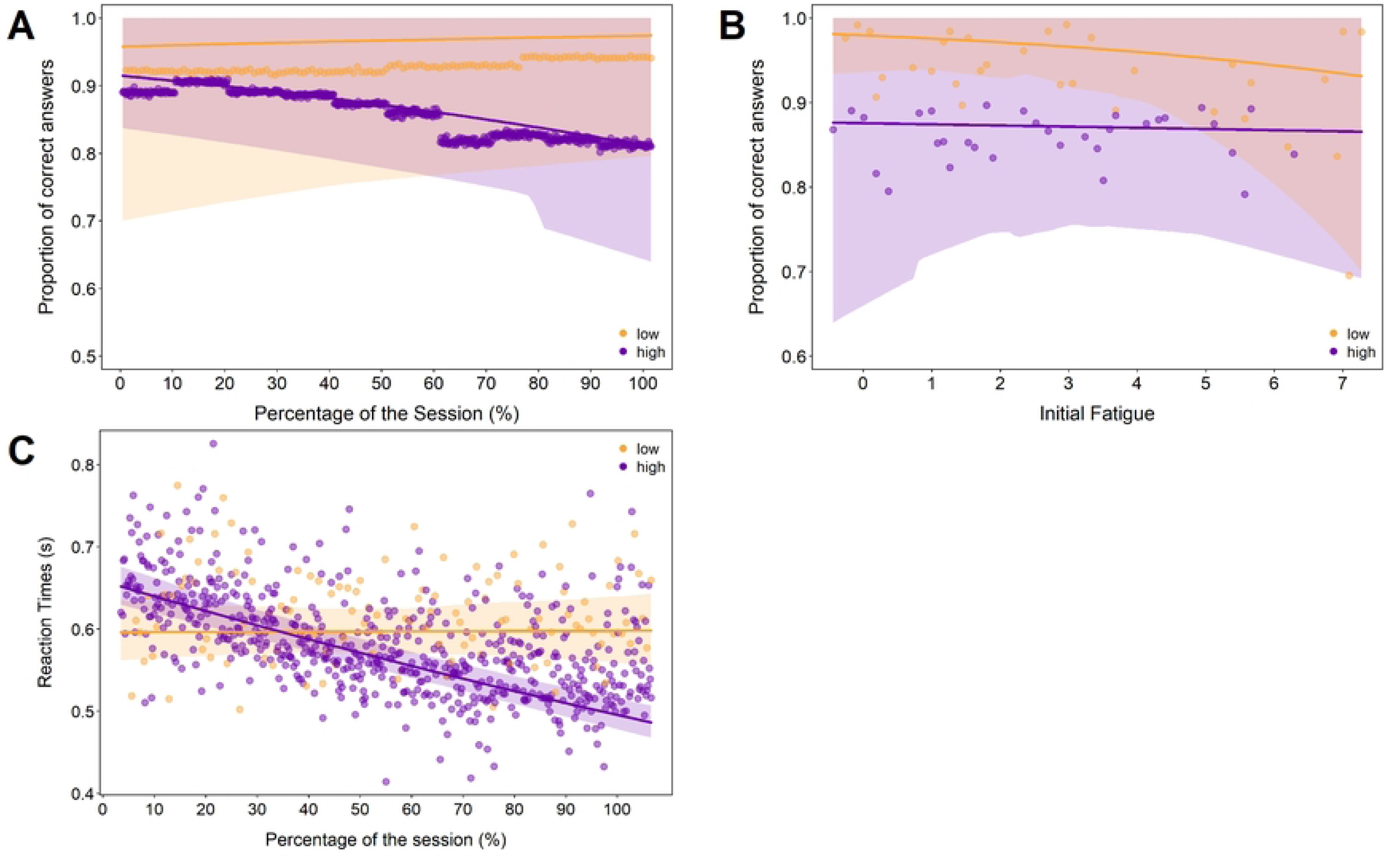
Performance in the cognitive demand tasks. *A*: Accuracy (proportion correct) over session progress (%). *B*: Accuracy as a function of initial fatigue. *C*: Reaction times (s) over session progress (%). The lines represent model fits for the low-demand (yellow) and high-demand (purple) condition; shaded areas represent the respective 95% confidence bands.

#### Effects of demand on pupil dynamics

For all pupil response models, interaction effects between demand condition and percentage of the session were significant. Across baseline, constriction and re-dilation - pupil diameter declined more strongly in the low-demand condition, whereas the high-demand task elicited more sustained pupil responses over time (*β* = 0.081, p <.001, semi-partial R² = .001, Figure 4A; *β* = 0.077, p < .001, semi-partial R² = .009, Figure 4B, and *β* = 0.072, p =.001, semi-partial R² = .007, Figure 4C respectively).

**Fig 4.**
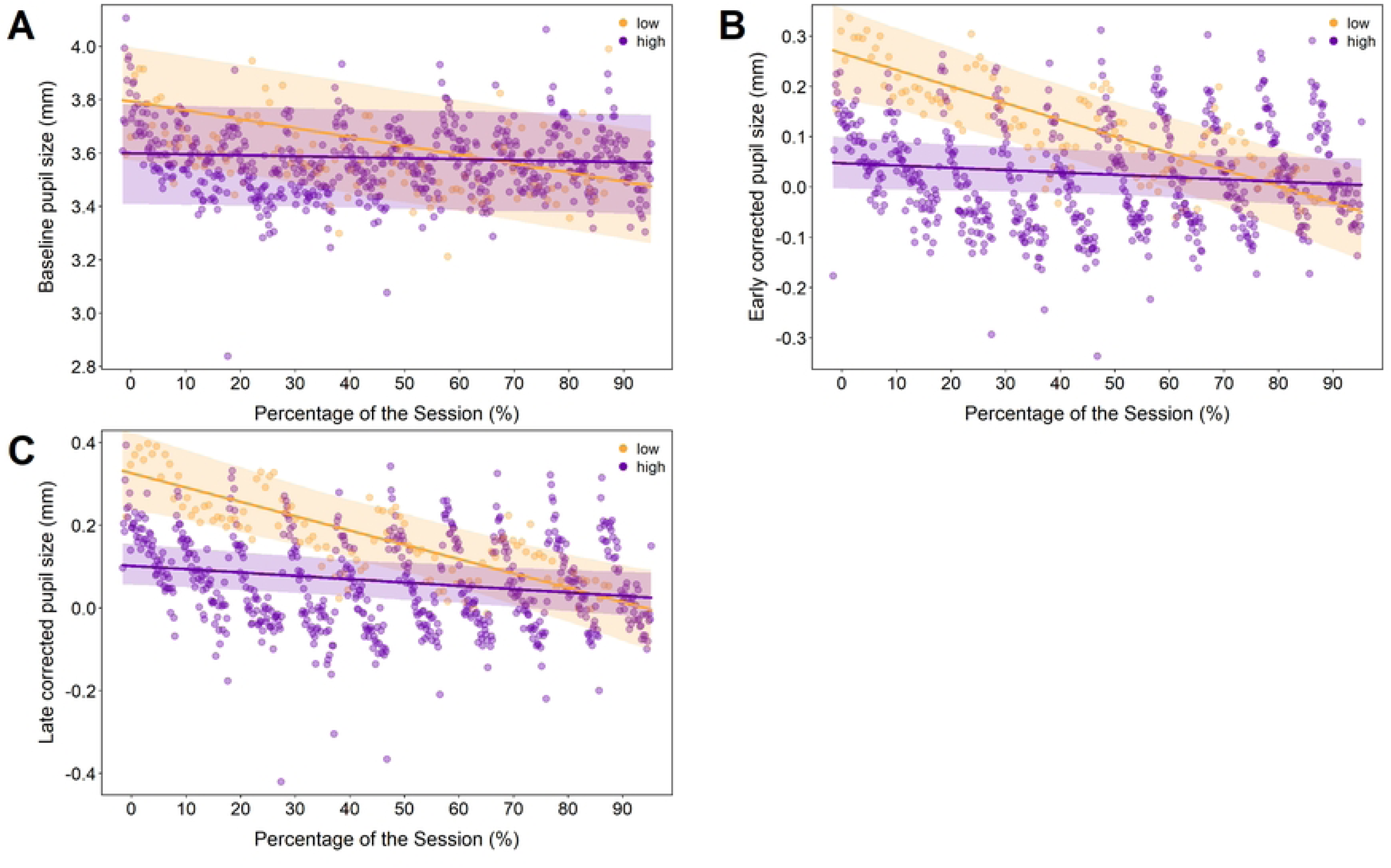
Pupil dynamics during the cognitive demand tasks. *A*: Baseline pupil diameter (mm). *B*: Early constriction diameter (baseline-corrected, mm). *C*: Late re-dilation diameter (baseline-corrected, mm), each shown over session progress (%). The lines represent model fits for the low-demand (yellow) and high-demand (purple) condition; shaded areas represent the respective 95% confidence bands.

### Motor-control task

We next tested whether prior demand condition affected motor performance and pupil dynamics in the motor-control task.

#### Effects of prior demand on motor-control performance

Across trials, RTs (object grab latency) decreased, indicating practice-related speeding independent of prior demand condition (*β* = -14.8, p = .002, semi-partial R² = .004; Figure 5A). Movement precision (deviation from the ideal trajectory) also improved with practice, again without a reliable difference between prior demand conditions (*β* = -0.034, p = .001, semi-partial R² = .004; Figure 5B). For movement duration, participants moved faster after the high-demand condition than after the low-demand condition and became faster over time (*β* = -0.045, p = .025, semi-partial R² = .004; Figure 5C and *β* = -0.012, p = .028, semi-partial R² = .001; Figure 5D respectively). For hit precision, performance tended to be higher after the low-demand condition and improved across trials (*β* = -0.97, p = .054; Figure 5E, and *β* = .06, p < .001; Figure 5F respectively).

**Fig 5.**
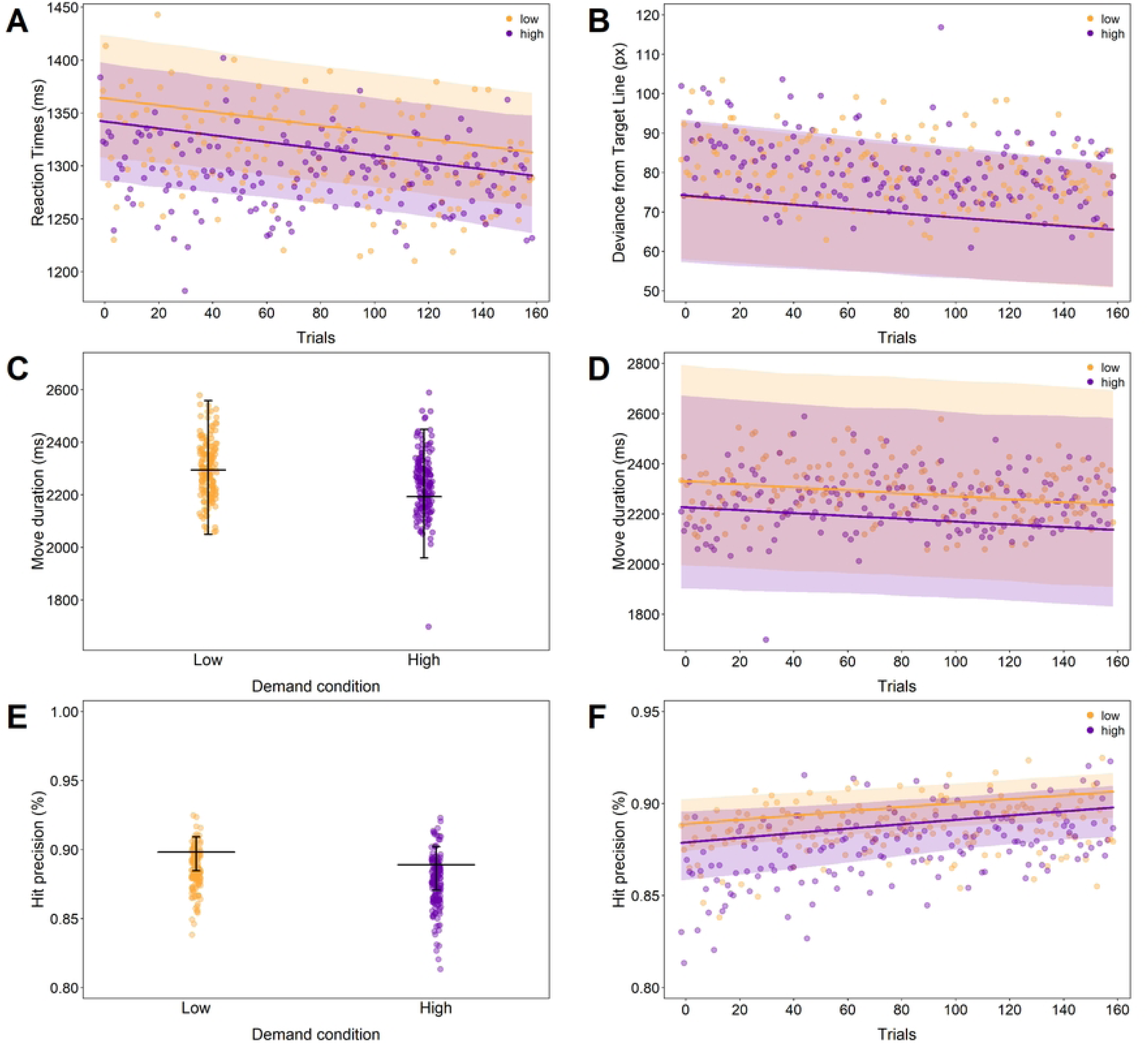
Performance in the motor-control task. *A*: RTs (ms) over trials. *B*: Movement precision as deviance from target line (px) over trials. *C–D*: Movement duration (ms) by demand condition and trials. *E–F*: Hit precision (%) by demand condition and trials. The lines represent model fits for the low-demand (yellow) and high-demand (purple) condition; shaded areas represent the respective 95% confidence bands. For plots by condition (C and E), horizontal lines represent model fits and vertical arrows represent the respective 95% confidence bands.

#### Effects of prior demand on pupil responses

Baseline pupil diameter during the motor-control task did not differ reliably between demand conditions and did not change systematically across trials. For early constriction (200-1200 ms), pupil responses did not vary significantly with trial number or demand condition alone, but there was a significant interaction with engagement: in the low-demand condition, higher engagement was associated with larger pupil responses, whereas in the high-demand condition, higher engagement corresponded to smaller responses (*β* = -0.078, p = .032, semi-partial R² = .025; Figure 6A). For late re-dilation (1200–2200 ms), a similar interaction with engagement emerged and re-dilation decreased modestly across trials (*β* = -0.082, p = .035, semi-partial R² = .022; Figure 6B, and *β* = -0.024, p < .001, semi-partial R² = .008; Figure 6C respectively). These findings indicate that the relationship between engagement and task-evoked pupil responses differed depending on prior cognitive demand.

**Fig 6.**
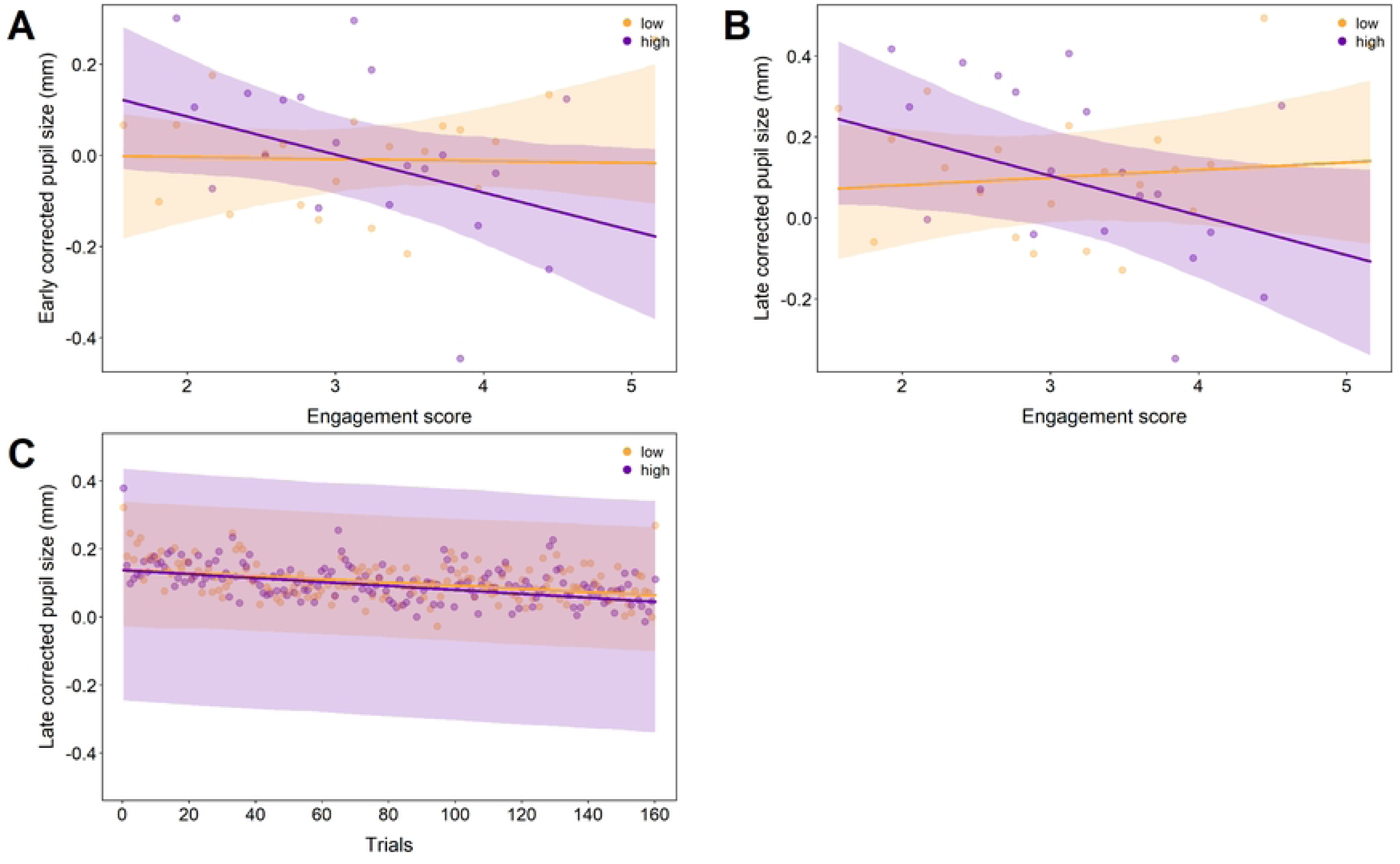
Pupil dynamics in the motor-control task. *A*: Early pupil constriction size (baseline-corrected, mm) over engagement. *B*: Late pupil re-dilation size (baseline-corrected, mm) over engagement. *C*: Late pupil re-dilation size (baseline-corrected, mm) over trials. The lines represent model fits for the low-demand (yellow) and high-demand (purple) condition; shaded areas represent the respective 95% confidence bands.

#### Performance predictions by pupil size

Finally, we examined whether trial-by-trial pupil metrics predicted performance in the motor-control task. Across all models, baseline pupil diameter did not reliably predict RT, movement precision, movement duration, or hit precision. Early constriction and late re-dilation were likewise unrelated to RT and movement precision. For movement duration, larger early constriction and greater late re-dilation were associated with slightly shorter movement durations *(β* = -0.013, p = .028, semi-partial R² = .001; Figure 7A, and *β* = -0.014, p = .038, semi-partial R² = .002; Figure 7B respectively). For hit precision, greater late re-dilation was linked to slightly lower spatial accuracy (*β* = -0.059, p = .001; Figure 7C). Effect sizes were small throughout; full statistics are reported in the Supplement.

**Fig 7.**
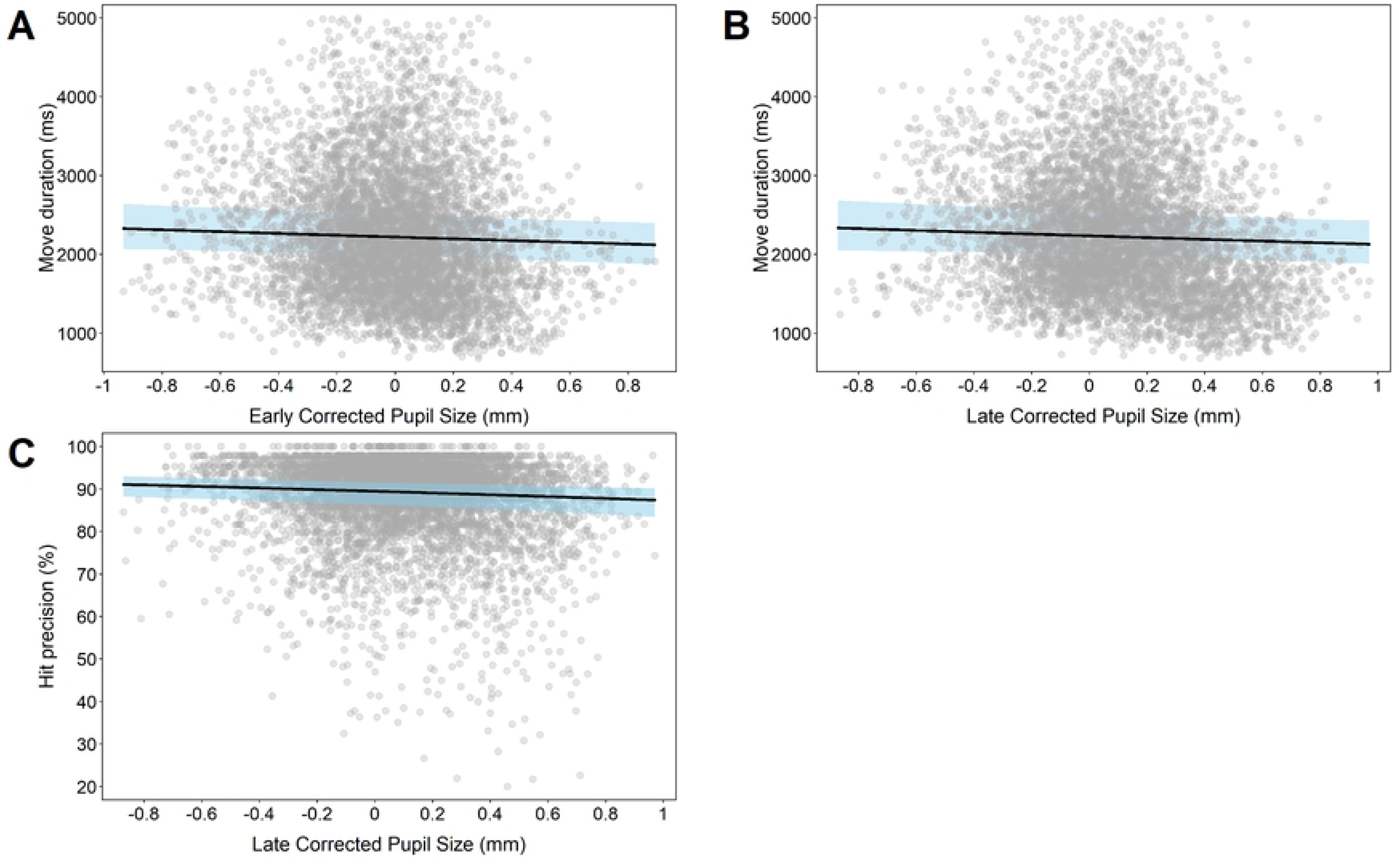
Pupil-performance relations in the motor-control task. *A–B*: Movement duration (ms) as a function of early pupil constriction and late pupil re-dilation size (baseline-corrected, mm). *C*: Hit precision (%) over late pupil re-dilation size (baseline-corrected, mm). Lines represent model fits; shaded areas represent 95% confidence bands.

## Discussion

This study examined how sustained cognitive demand shapes subsequent continuous motor control and tonic and phasic pupil dynamics in a controlled, HRI-inspired joystick task. Guided by the Compensatory Control Model (CCM) and the LC–NE adaptive gain framework, we examined whether pupil measures provide interpretable indicators of compensatory regulation during continuous motor control following sustained cognitive demand. This addresses the need for theory-guided interpretation of physiological signals alongside continuous motor behavior in sustained-demand contexts.

Three main findings emerged. First, the cognitive-demand manipulation produced robust behavioral and pupillary effects, validating the intended difference in sustained demand. Second, prior demand selectively influenced subsequent motor-control behavior, with participants adopting faster movement execution following higher demand, consistent with carry-over effects of compensatory regulation rather than uniform performance degradation. Third, phasic—but not tonic—pupil responses covaried with these behavioral adjustments. Importantly, these associations were statistically reliable but small in magnitude, constraining the strength of any performance-prediction claims.

H1 predicted that motor performance would differ as a function of prior cognitive demand. This hypothesis was only partially supported. Prior high demand selectively altered one motor-control metric: movements were faster following high demand, whereas endpoint accuracy showed only a non-significant trend toward reduced precision (p = .054). Other performance measures were unaffected. This pattern reflects a selective shift in the speed–precision balance rather than a uniform degradation of motor performance. In previous work, sustained cognitive fatigue has been associated with slower but more precise movements or with degraded fine motor control (55,56), a pattern opposite to the present findings. Importantly, both patterns are compatible with the Compensatory Control Model (CCM), which posits that operators can maintain overall task performance by reallocating effort and relaxing secondary goals when regulatory demands increase. Which specific performance components are prioritized seems to depend on task demands, incentives, and contextual constraints.

H2 predicted differences in tonic and phasic pupil responses following high versus moderate demand. Early constriction and late re-dilation were analyzed separately but showed mostly similar effects. Accordingly, we interpret both windows jointly as indicators of phasic engagement and compensatory regulation. Our hypothesis was only partly supported. Sustained demand robustly modulated pupil dynamics during the cognitive-demand task, but tonic baseline pupil diameter did not reliably differ between prior-demand conditions during the subsequent motor-control task. This pattern is consistent with evidence that baseline pupil diameter shows heterogeneous and task-dependent responses to sustained demand and is therefore better understood as reflecting broad arousal and motivational context rather than cognitive demand alone (31,33,57). In contrast, phasic pupil responses showed demand-related differences that depended on engagement. Within the CCM, sustained high demand is expected to recruit compensatory regulation, consistent with the robust demand-related pupil effects observed during the cognitive-demand task and with prior work linking pupil dynamics to effort regulation (26,31). The engagement-by-demand interaction in the motor task suggests that the physiological expression of engagement depended on prior demand: after low demand, higher engagement was associated with larger task-evoked responses, whereas after high demand, higher engagement was associated with smaller task-evoked responses. One interpretation is that following high demand participants entered the motor task in a more constrained regulatory state, such that additional engagement did not elicit increased phasic reactivity. Within LC–NE accounts, phasic pupil responses index momentary resource allocation and can be attenuated when responsivity is reduced or control costs accumulate (28,29,32). Crucially, these findings demonstrate that engagement does not map onto pupil size in a uniform way: depending on prior demand, higher engagement can be associated with either larger or smaller phasic pupil responses. For applied HRI, this underscores that pupil size cannot be interpreted as a context-free index of engagement or effort but must be evaluated in relation to prior task demands and regulatory state.

We also observed a decline in task-evoked pupil responses over time during the motor task. Similar time-on-task decreases have been reported previously and are often discussed in terms of fatigue or disengagement but have also been observed during motor learning and increasing task familiarity (1,12,31). In the present study, this decline coincided with behavioral improvement, which is compatible with a shift toward more efficient or stabilized regulation.

H3 predicted that phasic pupil responses would covary with fluctuations in motor performance and was weakly supported: larger late re-dilation—were associated with faster movement execution, and reduced endpoint precision. These associations indicate statistically detectable coupling between phasic pupil dynamics and movement execution, but effect sizes were very small (semi-partial R² ≈ .001–.002) limiting interpretability and precluding any strong performance-prediction claims. However, in early-warning or continuous-monitoring contexts, even weak but reliable physiological–behavioral couplings may be informative as one feature within multimodal operator-state models if replicated.

## Limitations

Although our joystick task introduced continuous, hand–eye coordination demands, it remains a laboratory simulation and does not capture the full complexity of real HRI scenarios. In more naturalistic settings, pupil dynamics will additionally be influenced by factors such as illumination changes, emotional cues, and multi-step collaboration. Further, the predictive relationships between pupil dynamics and motor performance were statistically reliable but extremely small. These effect sizes strongly limit theoretical inference and indicate that pupillometry alone is unlikely to support robust operator-state estimation.

## Practical implications

The present findings have several implications for the design of state-aware human–robot systems. First, performance metrics alone may be insufficient for detecting emerging operator strain: selective shifts in speed–precision balance occurred without uniform degradation, underscoring the value of continuous behavioral monitoring. Second, tonic pupil diameter should not be used as a standalone indicator of workload or fatigue, as it primarily reflects broad arousal and motivational context. In contrast, phasic, task-evoked pupil responses were more sensitive to regulatory dynamics, but only when interpreted relative to prior task demand and engagement. Finally, adaptive systems should avoid direct mappings from raw pupil size to operator state. Instead, pupillometry should be embedded within theory-guided, context-aware models that integrate behavioral dynamics and task history to reduce the risk of inappropriate or unsafe adaptations.

## Conclusions and future directions

This study examined how sustained cognitive demand shapes subsequent continuous motor control and pupil dynamics, using the Compensatory Control Model and LC–NE framework as interpretive anchors. The findings indicate that prior demand leads to selective behavioral adjustments rather than global performance decline and that phasic—but not tonic—pupil responses reflect engagement-dependent compensatory regulation. More broadly, the results reinforce that pupil dynamics do not index discrete states such as “fatigue” or “workload,” but reflect continuous, context-dependent regulation of control. Future research should extend this approach to more ecologically valid HRI settings, examine longer task sequences with repeated demand transitions, and evaluate whether combining pupil measures with behavioral and other physiological signals improves sensitivity to meaningful changes in operator regulation. Together, these findings provide an incremental but principled step toward theory-driven, context-aware use of pupillometry for operator-state modeling in adaptive human–robot systems.

## Acknowledgements

The authors thank the student research assistants who supported data collection for this study, as well as the participants for their time and participation.

## Supporting information

S1 Appendix. Model results.

